# Increased isoDGR motifs in plasma fibronectin are associated with atherosclerosis through facilitation of vascular fibrosis

**DOI:** 10.1101/2020.07.21.213397

**Authors:** Jung Eun Park, Xue Guo, Ken Cheng Kang Liou, Soe EinSi Lynn, Ser Sue Ng, Wei Meng, Su Chi Lim, Melvin Khee-Shing Leow, A Mark Richards, Daniel J Pennington, Neil E. McCarthy, Dominique P.V. de Kleijn, Vitaly Sorokin, Hee Hwa Ho, Siu Kwan Sze

**Affiliations:** School of Biological Sciences, Nanyang Technological University, 60 Nanyang Drive, Singapore 637551; Diabetes Center, Khoo Teck Puat Hospital, Singapore; Saw Swee Hock School of Public Health, National University of Singapore, Singapore; Cardiovascular and Metabolic Disorders Program, Duke-NUS Medical School, Singapore; Lee Kong Chian School of Medicine, NTU, Singapore; Department of Endocrinology, Tan Tock Seng Hospital, Singapore; Cardiovascular Research Institute, National University of Singapore, Singapore 119228; Christchurch Heart Institute, Department of Medicine, University of Otago, Christchurch 8140, New Zealand; Centre for Immunobiology, The Blizard Institute, Bart’s and The London School of Medicine and Dentistry, Queen Mary University of London, United Kingdom; Department of Vascular Surgery, UMC Utrecht, Utrecht University, Utrecht, the Netherlands; Netherlands Heart Institute, Utrecht, the Netherlands; Department of Cardiac, Thoracic and Vascular Surgery, National University Heart Centre, National University Health System, Singapore 119228; Department of Cardiology, Tan Tock Seng Hospital, 11 Jalan Tan Tock Seng, Singapore 308433

**Keywords:** Atherosclerosis, fibronectin, fibrillogenesis, isoDGR motif, integrin, deamidation

## Abstract

Abnormal matrix deposition on vessels and recruitment of inflammatory cells into the arterial wall are critical events in atherosclerotic plaque formation. Fibronectin protein is a key matrix component that exhibits high levels of deamidation in atherosclerotic plaques and blood plasma, but it is unclear how this structural change impacts on endothelial function or modifies interactions with recruited leukocytes. This study aimed to determine how deamidation-induced isoDGR motifs in fibronectin influence extracellular matrix accumulation on endothelial cells, and to investigate possible effects on integrin ‘outside-in’ signalling in matrix-bound monocytes which are key mediators of human atherosclerosis.

Blood plasma fibrinogen and fibronectin displayed marked accumulation of isoDGR motifs in ischemic heart disease (IHD) as determined by ELISA analysis of patients undergoing coronary artery bypass grafting compared with age-matched healthy controls. Biochemical and functional assays confirmed that isoDGR-containing fibronectin promoted activation of integrin β1 in monocytes and facilitated protein deposition and fibrillogenesis on endothelial cell layers. In addition, isoDGR interactions with integrins on the monocyte cell surface triggered an ERK:AP-1 signalling cascade that induced potent secretion of chemotactic mediators (including CCL2, CCL4, IL-8, and TNFα), that promoted further leukocyte recruitment to the assembling plaque.

Fibronectin deamidation forms isoDGR motifs that increase binding to β1 integrins on the surface of endothelial cells and monocytes. Subsequent activation of integrin ‘outside-in’ signalling pathways elicits a range of potent cytokines and chemokines that drive additional leukocyte recruitment to the developing atherosclerotic matrix and likely constitutes a key early event in progression to IHD.

## Introduction

Despite recent advances in therapeutic and preventive strategies, ischemic heart disease (IHD) remains a devastating disorder associated with extremely high morbidity and mortality across the world[3]. IHD comprises distinct pathologies termed ‘acute coronary syndrome’ and ‘chronic coronary artery disease’, which are routinely treated with a combination of medication, percutaneous interventions (often stent placements), and coronary artery bypass grafting (CABG) to circumvent occlusions. Early events in the pathogenesis of coronary artery disease (CAD) include endothelial dysfunction, vascular inflammation, and atherosclerotic plaque formation, ultimately leading to luminal narrowing, thrombotic occlusion of coronary arteries, and potentially acute myocardial infarction[2, 21]. While key processes of atherosclerotic plaque formation have been investigated in detail, it remains unclear how preceding events such as vascular fibrosis are initiated and which mechanisms are involved in later progression to clinically overt disease[26, 46].

Vascular fibrosis involves excess deposition of extracellular matrix (ECM) proteins such as collagen, proteoglycan and fibronectin in the arterial wall, leading to the reduced luminal diameter and increased vascular stiffness characteristic of atherosclerosis[25, 31, 32, 38]. Under normal conditions, ECM remodelling supports a variety of crucial repair processes including haemostasis, thrombosis, and angiogenesis[5, 17], but in a pathological context these events can become dysregulated and result in excessive ECM deposition and proliferation of vascular smooth muscle and endothelial cells[15, 43]. Fibronectin (FN) in particular binds to cell surface integrins via Arg-Gly-Asp (RGD) motifs, thereby facilitating adhesion of circulating monocytes to endothelial surfaces during appropriate inflammatory responses. However, this process is also strongly implicated in the vascular pathology of atherosclerosis[27]. Indeed, we have previously reported that extensive deamidation of NGR (Asn-Gly-Arg) sequences in ECM proteins results in ‘gain-of-function’ conformational switching to isoDGR (isoAsp-Gly-Arg) motifs[4, 7, 16, 20] that can mimic integrin-binding RGD ligands[11, 39]. Atypical isoDGR modifications were routinely detected in FN, laminin, tenascin C and many other ECM proteins derived from human carotid plaque tissues[4, 7, 16, 20], suggesting that these molecules may be capable of enhancing monocyte/macrophage binding to the atherosclerotic matrix via interactions with RGD integrin-binding pockets.

Long-lived ECM proteins are well recognized to accumulate degenerative proteinmodifications (DPMs), including spontaneous deamidation, under the influence of various microenvironmental stresses, flanking amino acid sequences, and genetic factors [18, 44]. DPMs are typically associated with loss of protein function and have long been regarded as critical events in human aging and implicated in various degenerative disorders. Importantly, recent data suggest that DPMs may also induce ‘gain of function’ structural changes that could play equally important roles in human pathology.[36, 41, 42]. Given our previous observation that FN is highly deamidated in atherosclerotic plaque tissues from CVD patients, we hypothesized that deamidation of soluble plasma FN may be a key early trigger for abnormal deposition, maturation, and stabilization of ECM components intrinsic to atherosclerotic matrix assembly. In the current study, we therefore aimed to determine the effect of deamidated fibronectin (containing isoDGR motifs) on ECM accumulation, matrix formation, and integrin ‘outside-in’ signalling in recruited leukocytes. We also investigated potential associations between ‘gain-of-function’ isoDGR-modified FN and the pathogenesis of IHD using blood plasma samples obtained from patients undergoing CABG. To do this, we generated a monoclonal antibody that enables specific detection of isoDGR epitope in damaged proteins via an enzyme-linked immunosorbent assay (ELISA). Using this approach, isoDGR motifs were found to be significantly enriched among plasma proteins derived from CABG patients, with particularly high enrichment being detected in both FN and fibrinogen. Subsequent biochemical and functional assays demonstrated that isoDGR-modified FN promoted activation of integrin β1 and facilitated protein deposition and fibrillogenesis on endothelial cell layers. In addition, interaction of isoDGR motifs with integrins on the surface of monocytes triggered an ERK:AP-1 signalling cascade that induced secretion of a range of potent chemotactic mediators that promoted further leukocyte infiltration of the assembling atherosclerotic matrix.

## Methods

### Patients and clinical samples

Study participants were recruited from patients undergoing coronary artery bypass grafting (CABG) or cardiac computed tomographic angiography (CCTA) in the Department of Cardiac, Thoracic and Vascular Surgery at the National University Heart Centre, National University Health System (NUHS) or the Department of Cardiology at Tan Tock Seng Hospital (TTSH). The study was approved by the institutional review boards of NTU (IRB-2017-01-013), NUHS (IRB-NUH-2009-0073) and TTSH (TTSH-2013-00930). Experimental procedures complied with the tenets of the Declaration of Helsinki. Informed written consent of each research subject was obtained prior to the inclusion in the study. For detailed demographic information and clinical history see; Table 1, Supplementary Tables 1 and 2, Fig. 1b. The control subjects were selected from patients who have been assessed by angiography on clinical suspicion and found to be non-CAD.

**Table 1.**
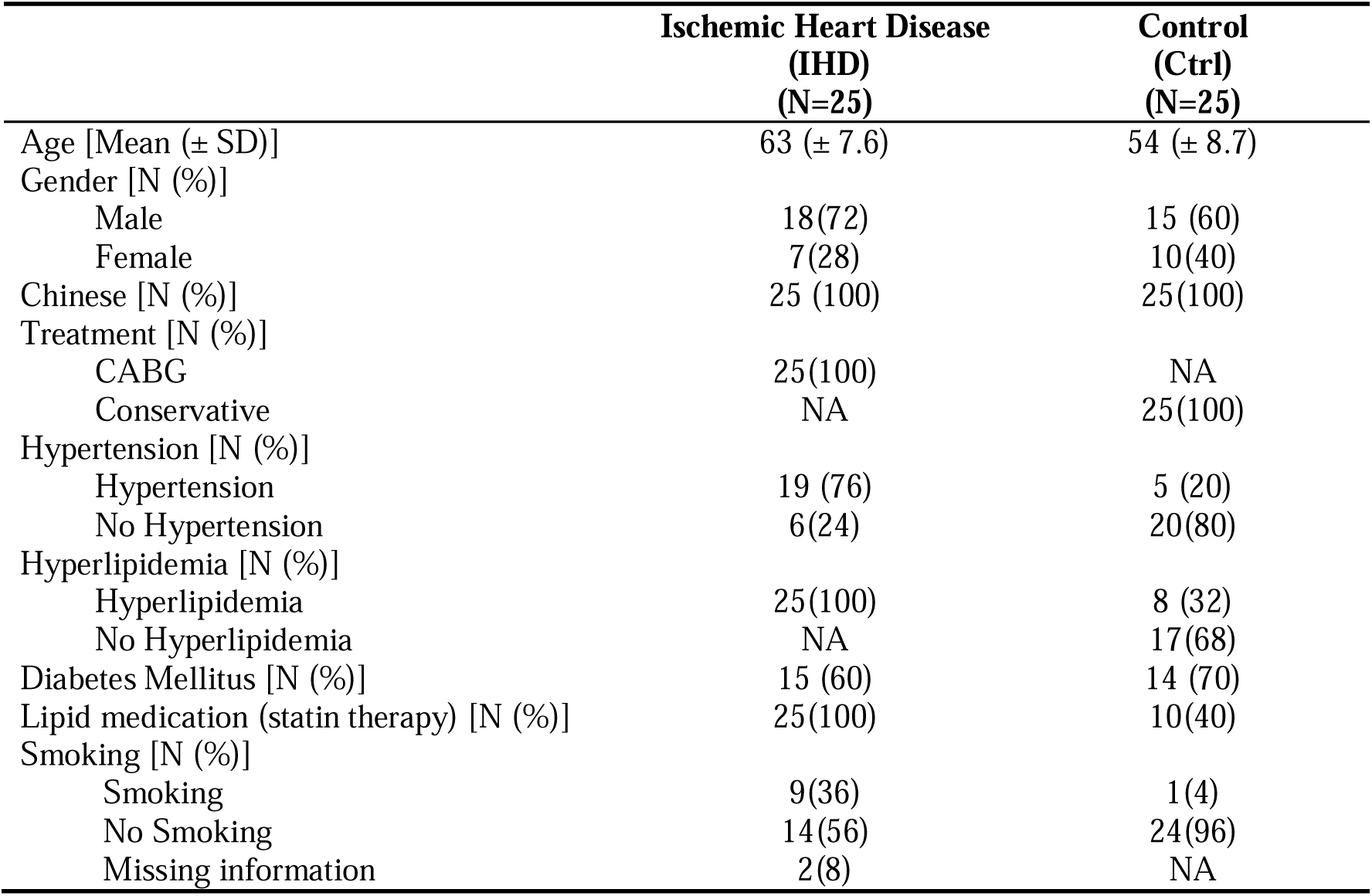
Demographic summary of clinical cohorts.

**Fig. 1.**
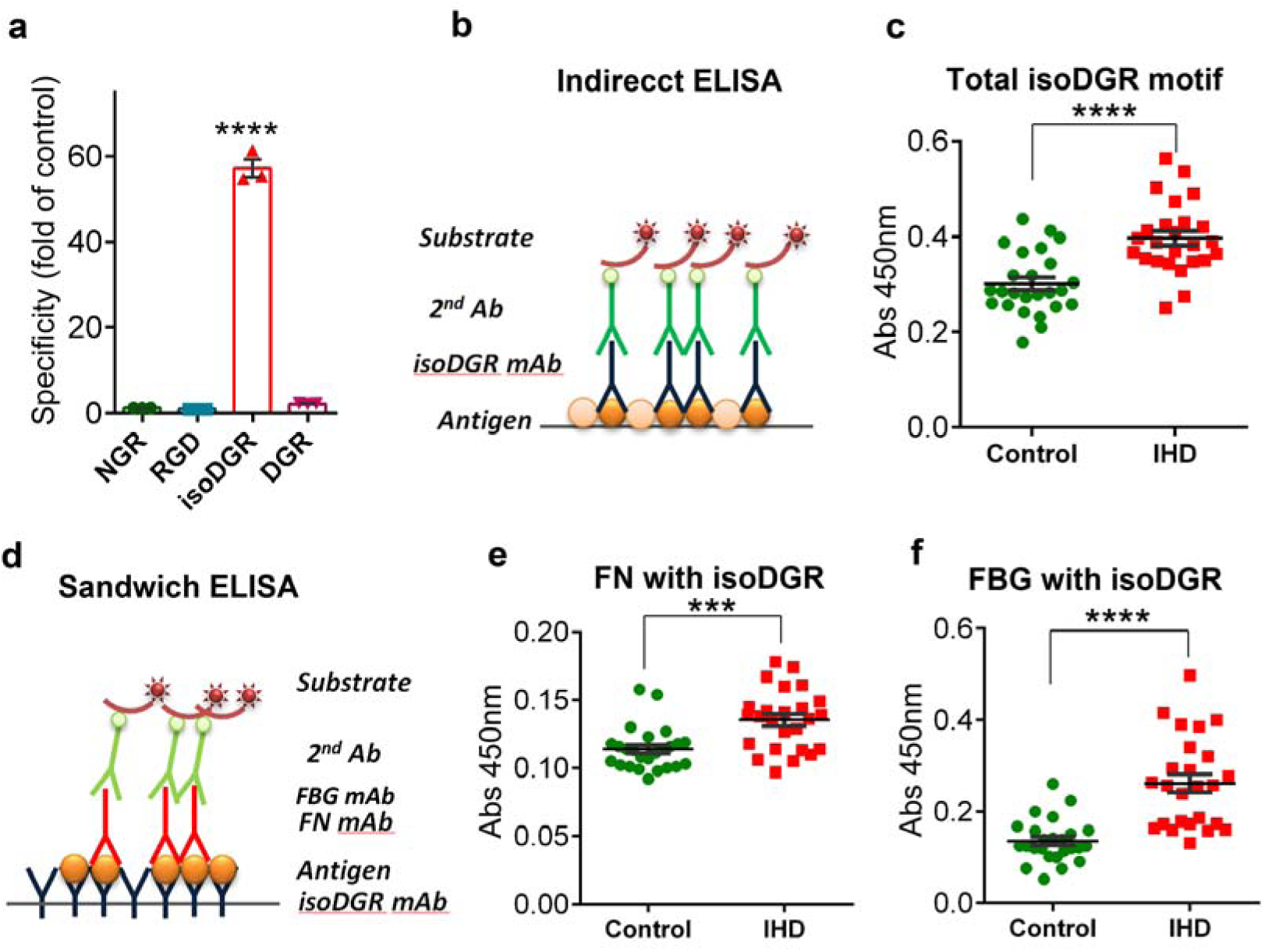
IsoDGR detection in blood plasma from patients with ischemic heart disease (IHD) **a** Monoclonal antibody specifically binds peptides that contain isoDGR motifs (but not those containing unmodified NGR, RGD, or DGR sequences); and **b** Schematic diagram of the indirect ELISA method. **c** Measurement of isoDGR levels in blood plasma samples from control donors (n=25) or IHD patients (n=25). **d** Schematic diagram of sandwich ELISA technique for measuring isoDGR motifs on fibronectin (FN) or Fibrinogen (FBG). **e** ELISA analysis of isoDGR motif level in plasma FN. **f** ELISA analysis of isoDGR motif level in plasma FBG. Plasma samples were from control donors (n=25) or IHD patients (n=25). ELISA assay was performed with three technical replicates (n=3). Statistical differences between groups were determined by unpaired T-test; *\*\*\*p=0*.*003, ****p<0*.*0001*.

### Cell lines and culture methods

Cell culture media and supplements were obtained from Life Technologies Holdings Pte Ltd. (Grand Island, NY, USA) unless otherwise specified. Human monocytic cell lines THP-1 and U937 were cultured in RPMI-1640 medium supplemented with 10% fetal bovine serum (FBS) and 1% penicillin/streptomycin (P/S) at 37°C in 5% CO_2_. Human umbilical vein endothelial cells (HUVECs) were cultured in DMEM containing 10% FBS and 1% P/S. All cell lines were tested for mycoplasma contamination by PCR according to published methods[49]. Before seeding onto fibronectin-coated 24 well plates, U937 and THP-1 cells were pre-stimulated with 100ng/ml Phorbol 12-myristate 13-acetate (PMA) (Sigma-Aldrich, St. Louis, MO, USA) for 1h in RPMI-1640 medium containing 1% FBS.

### Reagents, peptides, and antibodies

Human plasma fibronectin was purchased from Corning® (Corning, NY, USA). 5-Carboxyfluorescein N-Succinimidyl ester (CFSE) fluorescent dye was obtained from Sigma-Aldrich. AlexaFluor-488-microscale labelling kit was from ThermoFisher Scientific. The following peptides were produced by GL Biochem Ltd. (Shanghai, China); GC(isoD)GRCGG-(CH2-CH2-NH2), GCGRCGG-(CH2-CH2-NH2), GC(N)GRCGG-(CH2-CH2-NH2), and GCRGDCGG-(CH2-CH2-NH2). The molecular mass of each peptide was confirmed by MALDI-TOF mass spectrometry (MS) analysis. Antibodies against p44/42 MAPK (ERK1/2), phospho-p44/42 MAPK, and integrin β3 were purchased from Cell Signalling Technologies (Danvers, MA, USA). Antibodies against fibronectin, tubulin, and actin were from Santa Cruz Biotechnology (Santa Cruz, CA, USA) and anti-integrin β1 antibody (clone HUTS-4) was obtained from Chemicon (Merck Millipore, Billerica, MA). Rabbit polyclonal antibodies to fibronectin and fibrinogen were from Abcam (Cambridge, United Kingdom). SEAP reporter construct was purchased from BD Clontech (San Jose, CA, USA) and Phospha-Light™ was from Applied Biosystems (Bedford, MA).

### Generation of monoclonal antibody against isoDGR motifs

A custom-made mouse monoclonal antibody against the isoDGR motif was prepared by GenScript Corporation (Piscataway, NJ, USA). Briefly, two synthetic peptides, C(isoD)GRCK and (isoD)GR were conjugated with keyhole limpet hemocyanin (KLH) and used to immunize n=5 Balb/c mice. After cell fusion, ELISA screening against the antigen was used to select positive parental lines for sub-cloning and further expansion. We then used a combination of ELISA and dot blot assays to test reactivity of the selected hybridoma cells against four peptides containing either isoDGR or the dummy sequences DGR, RGD, and NGR. Based on the results of these assays, we selected a single hybridoma that displayed high specificity for the isoDGR motif but lacked reactivity to DGR, RGD, or NGR (Fig. 1a). Monoclonal antibody was purified from the supernatant of hybridoma cell cultures using protein A/G agarose columns according to the manufacturer’s instructions (Thermo Fisher Scientific, San Jose, USA).

### Indirect and sandwich ELISA for isoDGR detection in human plasma samples

To detect total deamidated NGR motifs in plasma samples, a total of 5µg plasma protein in 100µl PBS was loaded into 96-well plate (Maxisorp, NUNC) and incubated at 4°C overnight to coat the wells. The plate was then blocked with 3% BSA in PBS for 1h at room temperature (RT) and washed twice with PBS containing 0.05% Tween 20 (PBST). Mouse monoclonal anti-isoDGR antibody (5µg/ml) was subsequently added to each well and incubated for 2h at RT. The secondary anti-mouse IgG antibody conjugated to horseradish peroxidase (Pierce, Rockford, IL) was diluted 1:5000 in blocking solution then added to each well and incubated for 1h at RT. The plate was rinsed four times with PBST before adding 100μl Turbo-TMB (Thermo Scientific) to each well and incubating for 10min at RT. The reaction was stopped by adding 100μl of 2N H_2_SO_4_ before measuring absorbance at 450nm (TECAN microplate reader, Maennedorf, Switzerland).

To enable rapid detection of isoDGR motifs in fibronectin and fibrinogen from human plasma samples, we next developed a sandwich ELISA approach using our custom antibodies. A total volume of 100µl anti-isoDGR monoclonal antibody was added to each well for overnight incubation at 4°C, after which the wells were washed three times with 200µl PBS. Non-specific binding sites were blocked using 3% BSA in PBS for 1h at RT. After washing, 100µl samples were added to each well and incubated for 2h at RT. Following washing steps, rabbit polyclonal antibodies against fibronectin or fibrinogen were added and incubated for 2h at RT. After washing with PBST, the HRP-coupled donkey monoclonal anti-rabbit antibodies (Cell Signalling) was added for 1hr at RT. Next, the wells were washed 3 times with PBST and 100µl TMB was added to each well for 10min to enable colour development. To stop the colour reaction, 100µl 2N H_2_SO_4_ was added before reading optical density at 450nm.

### Fibronectin deamidation and tandem mass spectrometry

Two forms of plasma FN (native and deamidated) were used in the experiments. Native fibronectin was dissolved in ammonium acetate buffer at pH 6.5, while deamidated FN was prepared by incubating FN in 0.1M ammonium bicarbonate buffer (pH 8.0) for 48h at 37°C before adjusting to pH 6.5 by addition of 1% acetic acid. Native and deamidated FN structures were confirmed by LC-MS/MS and then used for cell treatment or plate coating.

### In-solution proteomic sample preparation

A 20μg mass of fibronectin protein in solution (pH 6.5) was reduced with 10mM dithiothreitol for 2h at RT, then alkylated using 55mM iodoacetamide for 1h at RT in the dark. The sample was then incubated at 37°C for overnight trypsin digestion (trypsin 1:50 protein w/w ratio; Promega, Madison, WI). The digestion reaction was quenched by adding formic acid (FA) until the final acid concentration reached 0.5%. Tryptic peptides were dried in a vacuum concentrator (Concentrator Plus, Eppendorf AG, Hamburg, Germany).

### Label-free LC-MS/MS analysis

The dried peptides were reconstituted in 3% acetonitrile, 0.1% FA buffer followed by vortexing and sonication for 1h. A 0.2μg quantity of the sample was then injected into a Dionex Ultimate 3000 RSLCnano system coupled to a Q Exactive instrument (Thermo Fisher Scientific, Inc.). Peptide separation was performed on a Dionex EASY-Spray 75 μm x 10 cm column packed with PepMap C18 3μm, 100 Å (Thermo Fisher Scientific, Inc). Sample peptides were first loaded into an acclaim peptide trap column via the autosampler. Separation of peptides was performed with mobile phase solvent A (0.1% FA in HPLC water) and solvent B (0.1% FA in 90% ACN) at flow rate of 300nl/min with a 60min gradient as follows: 3−30% B for 45min, 30−50% B for 9min, 50−60% B for 1min, 60% B for 2min, and finally isocratic at 3% B for 3min. The sample was sprayed with an EASY nanospray source (ThermoFisher Scientific, Inc.) at an electrospray potential of 1.5 kV. The Q Exactive was set to perform data acquisition in positive ion mode and a full MS scan (350-1,600 m/z range) was acquired at a resolution of 70,000 at m/z 200. The 10 most intense ions were selected for higher energy collisional dissociation (HCD) fragmentation using 28% normalized collision energy. The AGC setting was 1E6 for the full MS scan and 2E5 for the MS2 scan. Single and unassigned charged ions were excluded from MS2.

### Label-free proteomic data analysis

Raw data files were converted into the mascot generic file format (MGF) using Proteome Discoverer version 1.4 (Thermo Electron, Bremen, Germany) with the MS2 spectrum processor for de-isotoping and de-convoluting MS/MS spectra. Label-free raw data file searches were carried out using an in-house Mascot server (version 2.6.02, Matrix Science, MA) with precursor ion tolerance of 10ppm and fragment ion tolerance of 30ppm. The UniProt human database was used for protein database searches (downloaded on June 23, 2015, including 180,822 sequences and 71,773,890 residues). Variable modifications were set as deamidation of N and Q, as well as oxidation of M, while carbamidomethylation of C was set as a fixed modification. Two missed trypsin cleavage sites per peptide were tolerated.

### FN fibrillogenesis and DOC solubility assay

Native and deamidated FN were labelled with Alexa488 dye and used for exogenous plasma fibronectin assembly assays. Briefly, HUVECs were seeded onto coverslips and allowed to adhere and grow to 90-100% confluency. Subsequently, cells were treated with either native or deamidated FN (25µg/ml) in 2% FBS-DMEM media containing PMA (100ng/ml) and incubated for the indicated times at 37°C with or without steady flow (100rpm). FN deposition and/or matrix assembly were monitored using fluorescence microscopy and subsequently processed for sodium deoxycholate (DOC) solubility assays as previously described[47]. Briefly, HUVECs were put on ice and lysed with cold DOC lysis buffer (2% DOC; 20mM Tris–HCl, pH 8.8; 2mM PMSF; 2mM EDTA; 2mM iodoacetic acid; 2mM N-ethylmaleimide). Cells were lysed using a 26G syringe needle and immediately centrifuged at 15,000rpm for 15min at 4°C. The supernatant was collected as DOC soluble fraction while the pellet was washed with DOC buffer and solubilized in 25μl SDS buffer (1% SDS; 20mM Tris–HCl, pH 8.8; 2mM PMSF; 2mM EDTA; 2mM iodoacetic acid; 2mM N-ethylmaleimide). DOC-soluble and DOC-insoluble fractions were subsequently resolved by SDS–PAGE using 7.5% polyacrylamide gels, then transferred onto nitrocellulose membranes and immunoblotted with anti-FN mAb.

### Quantitative real-time PCR

Quantitative real-time PCR (qRT-PCR) was used to measure mRNA expression levels of cytokines by U937 and THP-1 monocytic cells. Total RNA extraction was performed using Nucleospin RNA kits (MACHEREY-NAGEL GmbH & Co.) according to the manufacturer’s protocol. qRT-PCR was performed using a CFX96 Real-Time PCR Detection System (Bio-Rad) with KAPA SYBR® FAST qPCR Master Mix. To normalize quantification, levels of 18s RNA detected on the same plate as the target genes were used as internal controls. The qRT-PCR reaction was as follows; denaturation at 95°C for 15s followed by annealing at 60°C for 15s and extension at 72°C for 15s then final extension at 72°C for 5min over a total of 40 cycles. The primer sequences of target genes were as follows: CCL4 F 5’-AAGCTCTGCGTGACTGTCCT-3’, CCL4 R 5’-GCTTGCTTCTTTTGGTTTGG-3’; MMP9 F 5’-TTGACAGCGACAAGAAGTGG-3’, MMP9 R 5’-GCCATTCACGTCGTCCTTAT-3’; CCL2 F 5’-CAATCAATGCCCCAGTCACC-3’, CCL2 R 5’-TCGGAGTTTGGGTTTGCTTG-3’; TNFα F 5’-GTCAACCTCCTCTCTGCCAT-3’, TNFα R 5’-CCAAAGTAGACCTGCCCAGA-3’; IL-8 F 5’-CAGTTTTGCCAAGGAGTGCT-3’, IL-8 R 5’-ACTTCTCCACAACCCTCTGC-3’; 18s RNA F 5’-GTAACCCGTTGAACC CCATT-3’, 18s RNA R 5’-CCATCCAATCGGTAGTAGCG-3’.

### Transwell migration assay

Native or deamidated FN were coated onto 96-well plates (2.5µg/100µl volume per well) via overnight incubation at 4°C. Control wells were prepared with 1% bovine serum albumin (BSA) in PBS and transmigration assays were carried out using n=5 technical replicates. Briefly, U937 cells were primed with PMA (100ng/ml) for 1h at 37°C then re-suspended in 1% FBS-RPMI medium and seeded onto FN-coated plates at a density of 1 × 10^5^ cells/well for 6h. Thereafter, 4 × 10^4^ CSFE-labelled, non-activated U937 cells were added to matrigel-coated inserts and incubated for a further 24h at 37°C. Cell transmigration in response to molecules secreted by FN-bound monocytes was monitored using fluorescence microscopy and quantified by crystal violet staining.

### Secreted alkaline phosphatase (SEAP) reporter assay

U937 cells were reverse-transfected with 50ng AP1-SEAP or NFκB-SEAP vector, then seeded onto native or deamidated FN-coated 96-well plates and incubated for 24h. Culture supernatants were collected, heated at 65°C for 30□min, then assayed for alkaline phosphatase activity as follows; 30µl of supernatant was incubated with 120µl assay buffer for 5min, after which 1:20 diluted CSPD substrate was added and the samples were read on a TECAN microplate reader (Maennedorf, Switzerland).

### Western blot analysis

Cells were washed in ice-cold PBS and lysed in modified RIPA buffer (50mM Tris–HCl, 150mM NaCl, 1% NP-40, pH 8.0, 1 × protease inhibitor cocktail, phosphatase inhibitors). Lysates were clarified by centrifugation (16,000□×□g, 30min) and subjected to western blotting using the indicated primary antibodies at 1:1000 dilution. Protein-antibody conjugates were visualized using a chemiluminescence detection kit (Thermo Fisher Scientific).

### Statistical analysis

Statistical analyses were performed using GraphPad Prism 5.0 (GraphPad Software, Inc., San Diego, CA). Differences between groups were assessed by Student’s t-test and p<0.05 was considered significant.

## Results

### IsoDGR motifs are increased in blood plasma from patients with cardiovascular disease (CVD)

Deamidation of NGR (Asn-Gly-Arg) amino acid sequences in extracellular matrix (ECM) proteins creates ‘gain-of-function’ isoDGR motifs (isoAsp-Gly-Arg) implicated in atherosclerotic plaque formation, so we developed a specific monoclonal antibody against isoDGR to enable analysis of this structure in human patient samples (Fig. 1a). Blood plasma from 25 patients with ischemic heart disease (IHD) undergoing coronary artery bypass grafting (CABG) were screened by ELISA for total levels of isoDGR motifs and compared with 25 controls lacking any history of hypertension or hyperlipidemia (Table 1, Fig. 1b, Supplementary Tables 1 and 2). As shown in Fig. 1c, levels of isoDGR motifs were significantly increased in IHD patients, suggesting increased incidence of NGR deamidation relative to control plasma donors (p<0.0001). In previous analyses of human carotid plaque tissues, we identified high levels of deamidation in core proteins fibrinogen (FBG) and fibronectin (FN) which are known to play critical roles in matrix assembly and thrombosis[9, 33]. Using the isoDGR-specific mAb generated in the current study, we next developed a sandwich ELISA protocol to facilitate analysis of deamidation levels in blood plasma FN and FBG obtained from patients with IHD (Fig. 1d). These data confirmed that FN and FBG in blood plasma from IHD patients exhibit significantly higher levels of isoDGR motifs than do the same plasma proteins from control donors, thus implicating the deamidated variants in the pathological progression of CVD (Fig. 1e, 1f).

### Endothelial cell exposure to deamidated fibronectin (FN) increases activation of integrin β1

We next sought to determine whether deamidated FN can enhance integrin activation on endothelial cells. When incubated under alkaline conditions (0.1M ammonium bicarbonate, pH 8.5), the NGR motif of FN protein is susceptible to non-enzymatic deamidation at the Asn residue which undergoes conversion to isoAsp (confirmed here using LC-MS/MS analysis; Fig. 2A and B). The new isoDGR motif is located on FN subunit I (red color) and faces outwards from the main body of the molecule, suggesting that this *de novo* structure is likely to be readily accessible for integrin binding. To test this, we added either native or deamidated FN (25µg/ml) to PMA-primed human umbilical vein endothelial cells (HUVECs) for 24h and performed immunofluorescent staining for active integrin β1 (using antibody clone HUTS-4 which recognizes epitopes in the 355–425 region of active β1 subunits[29]). Levels of activated integrin β1 where markedly increased on HUVECs exposed to deamidated FN relative to those treated with the native protein alone, and active integrin β1 was more clearly distributed to the plasma membrane in the presence of FN featuring isoDGR motifs (Fig. 2c, 2d). To further elucidate activation signals triggered by modified FN, we next exposed PMA-primed HUVECs to a range of synthetic peptides containing either NGR, RGD, or isoDGR motifs for 24h before repeating the immunofluorescent analyses. Unlike peptides containing NGR or RGD sequences, ligands that featured an isoDGR motif potently induced integrin β1 activation and distribution to the membrane region of PMA-primed HUVECs (Fig 2E). However, HUVECs routinely failed to react to FN in the absence of PMA priming, suggesting that basal integrin activation is prerequisite for isoDGR motifs to influence endothelial β1 function.

**Fig. 2.**
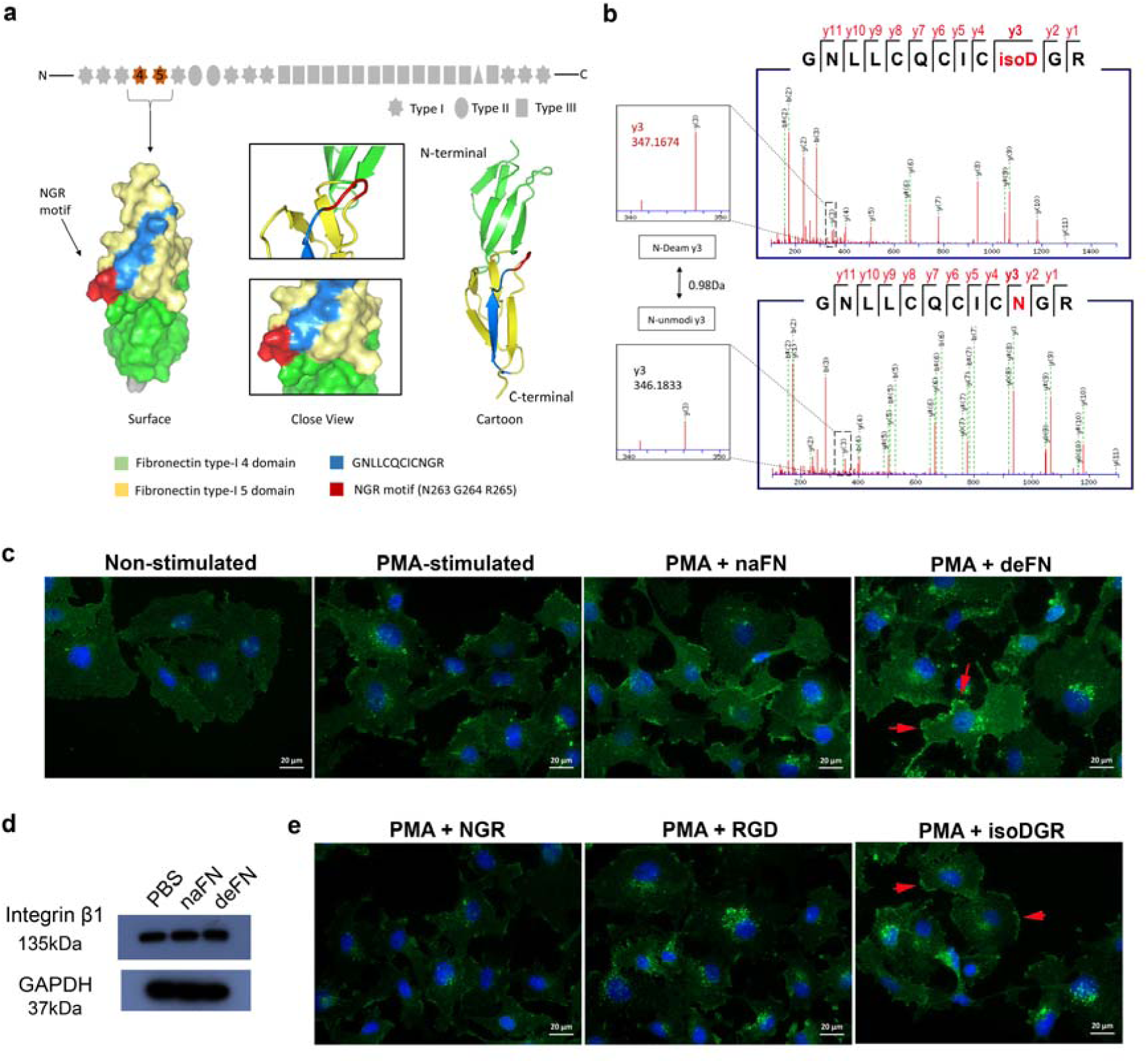
IsoDGR-containing fibronectin enhances integrin β1 activation on human endothelial cells. Structural schematic and tandem mass spectrometry confirmation of fibronectin (FN) modification at residue Asn263 following protein deamidation *in vitro*. **a** The NGR motif (red) is located within the FN type I domain and faces outwards from the main body of the molecule. **b** CID mass spectrum of peptide _254_GNLLCQCICNGR_265_ confirming deamidation of the Asn residue (+0.98Da) accompanied by characteristic y ion series (y2–y11, solid red lines), and neutral losses (dashed green lines) of NHCO (43 Da) and NH3 (17 Da). **c** HUVEC endothelial cells treated with deamidated-FN (25µg/ml) displayed higher levels of integrin β1 activation and distribution to the plasma membrane (red arrows) than were observed among control cells exposed to the native protein (n=3, scale bar 20 µm). **d** Total expression levels of integrin β1 were comparable between conditions upon assessment by western blot (n=3). **e** Activation of integrin β1 in these assays required the presence of peptides containing isoDGR motifs but not the alternative sequences NGR or RGD. Key: Native fibronectin, naFN; deamidated fibronectin, deFN (n=3, scale bar 20 µm).

### Fibronectin NGR-to-isoDGR transition facilitates fibrillogenesis and insoluble matrix formation

FN:integrin interactions initiate step-wise conformational changes in FN-FN interactions on cell surfaces that support conversion to insoluble fibrils / fibrillogenesis[37, 48]. Since our data suggested that deamidated FN can enhance activation of integrins – which are the principal FN receptors in ECs – we next examined whether protein deamidation was able to modify normal processes of FN polymerization, accumulation, and fibril formation. To do this, we incubated PMA-primed HUVECs for variable duration with either native or deamidated plasma FN (25µg/ml) that had been pre-labelled with AlexaFluor488 fluorescent dye. When visualized by microscopy, we rarely observed any deposition of native FN on HUVECs after 6h exposure, and FN aggregation could only be detected after an extended 18h incubation period (Fig. 3a). In contrast, deposition of deamidated FN was clearly visible within just 6h and FN aggregates on HUVECs were greatly increased over time. Since vascular endothelial cells are exposed to constant mechanical force *in vivo*,[8] we next investigated the dynamics of native versus deamidated FN assembly on PMA-primed HUVECs subjected to continuous flow (CO_2_ shaking incubator for 18h duration). Under these conditions, deposits of deamidated FN converted to stretched and unfolded forms on endothelial cell surfaces, whereas native FN accumulated in greater quantities but failed to form extended fibrils, suggesting that deamidation of plasma FN facilitates matrix assembly and formation of dense fibril networks (Fig. 3b). To further test whether isoDGR-containing FN can accelerate the formation of stable and mature fibrillary networks, we next used a deoxycholate (DOC) detergent assay to assess how deamidated FN impacts ECM solubility, which is a critical determinant of overall tissue stability, rigidity, and shape[19, 30]. Data supported the microscopic observation of isoDGR-mediated FN fibrillogenesis in a quantitative manner. Western blot analysis of proteins obtained by DOC buffer-extraction from HUVECs revealed a significant increase in insoluble FN content among cells that had been exposed to isoDGR-containing variant relative to the unmodified protein (p=0.0185; Fig. 3c).

**Fig. 3.**
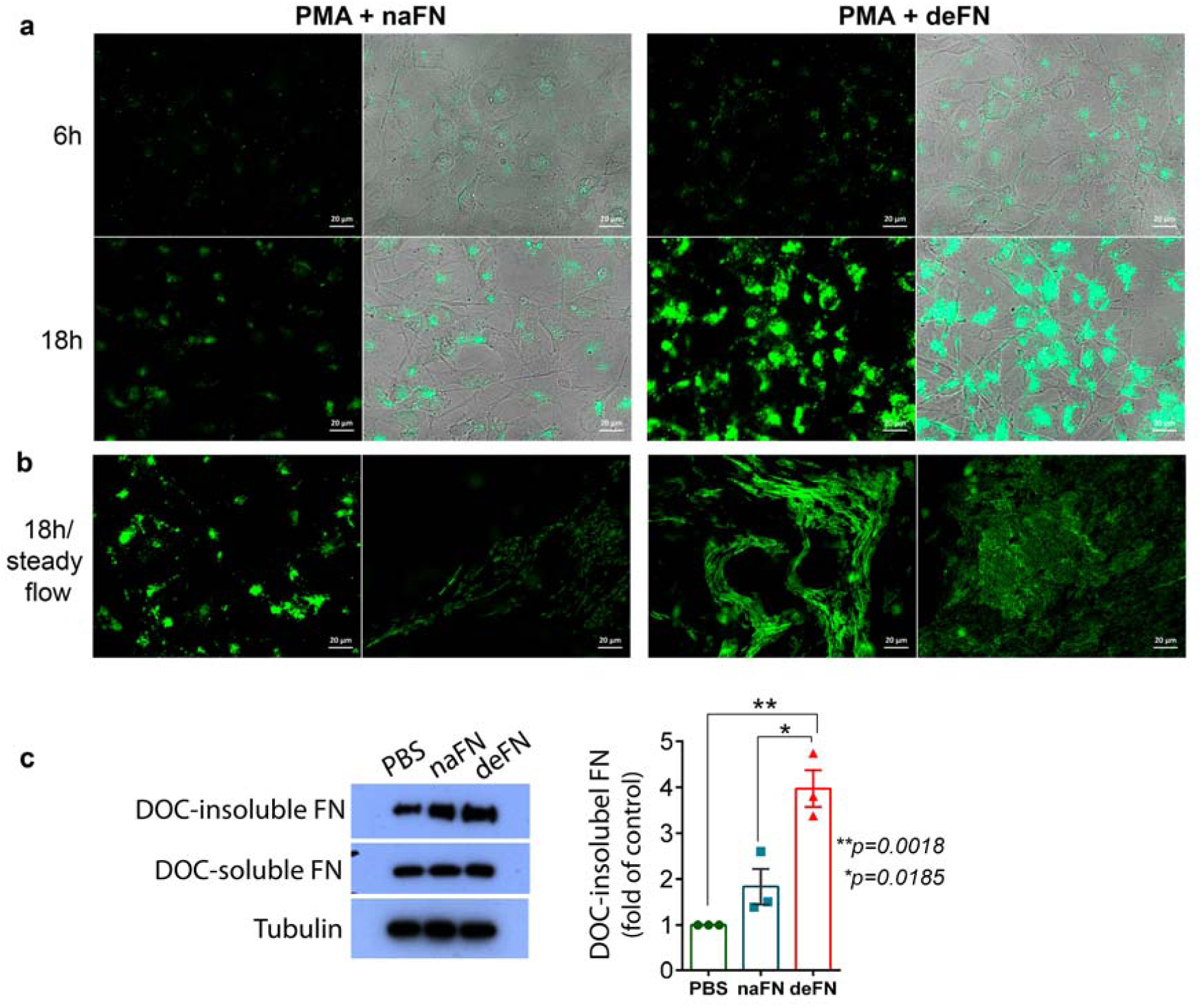
IsoDGR-containing fibronectin accelerates protein fibrillogenesis on endothelial cells. **a** PMA-primed HUVECs were incubated for 6h or 18h in the presence/absence of either native or deamidated fibronectin (FN) that had been pre-stained with Alexa Fluor 488 dye. Accumulation of exogenous FN on endothelial cell surfaces was markedly increased by FN deamidation (n=3, scale bar 20 µm). **b** When the same experiment was repeated under conditions of steady flow (100 rpm for 18h duration), deamidated FN was observed to extend and form stretched fibrils associated with matrix assembly, whereas native FN displayed only limited accumulation on the HUVEC monolayer (n=3, scale bar 20 µm). **C** Upon completion of the steady-flow assay, HUVECs were treated with 1% DOC detergent to assess matrix solubility, which revealed that deamidated FN strongly resisted DOC lysis, consistent with reduced solubility of the isoDGR-modified fibrils (n=3 biological replicates). Quantification of western blot images was by Image J software. Statistical differences between groups were determined by unpaired T-test; *p=0.015, **p=0.0018.

### IsoDGR formation induces activation of integrin ‘outside-in’ signaling pathways

In previous work, we reported that monocyte adhesion to FN-coated plates can be increased by deamidation of NGR motifs, and that this interaction is primarily mediated by cell-surface integrins.[16] Consequently, in the current study we sought to determine the biological consequences of integrin-mediated monocyte adhesion to isoDGR-containing FN protein. To this end, PMA-primed U937 monocytic cells were cultured in 24-well plates coated with either native or deamidated FN. After 24h incubation, the cells were harvested, subjected to total protein extraction, and western blot analysis was performed to identify expression levels of molecules including integrin β3 and downstream signaling molecules such as AKT and mitogen-activated protein kinase (MAPK)/ERK. As shown in Fig. 4a, PMA-treated U937 cells displayed high basal levels of integrin β3 expression that were marginally increased upon adhesion to deaminated FN, whereas AKT phosphorylation at Ser473 was suppressed upon PMA exposure, irrespective of FN binding state. Instead, interaction of monocyte integrins with deamidated FN appeared to modulate the alternative MAPK-ERK signaling pathway, since levels of phosphorylated ERK1/2 were increased among monocytes exposed to deamidated FN during culture. To further investigate the effects of NGR-to-isoDGR molecular switching on downstream pathways of integrin ‘outside-in’ signalling, we next used reporter constructs containing the secreted alkaline phosphatase (SEAP) gene under the control of either AP-1 or NF-κB promoters to quantify pathway activation. Monocytes binding to deamidated FN displayed a >2-fold increase in AP-1-induced SEAP activity compared to native FN-exposed cells, indicating that isoDGR motifs can promote signal transduction through this axis and potentially upregulate downstream gene targets of AP-1. Intriguingly, isoDGR-triggered integrin signalling appeared independent of NF-κB which was not activated in our assays despite this pathway playing a central role in regulating inflammation[28].

**Fig. 4.**
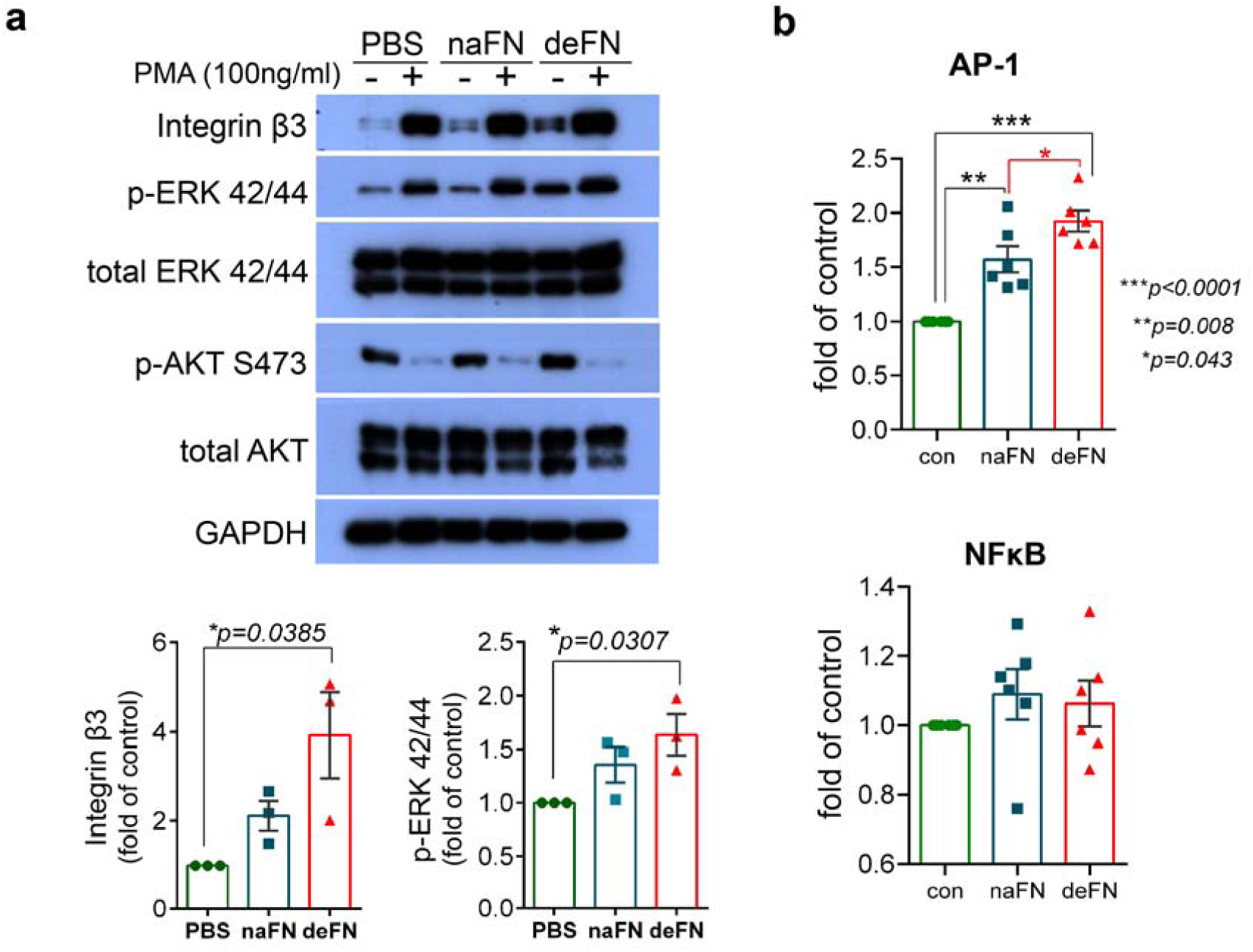
Monocyte adhesion to isoDGR-containing fibronectin triggers integrin ‘outside-in’ signalling. **a** PMA-primed U937 monocytes were cultured in 24-well plates coated with either native or deamidated FN for 24h, during which time ERK phosphorylation was induced only in cells exposed to the isoDGR-modified protein (lower panel). Quantification of western blot images was performed using Image J software and compared against GAPDH protein loading control (n=3). Statistical differences between groups were determined by unpaired T-test. **b** U937 cells were reverse-transfected with AP-1 or NFκB-SEAP vector and cultured on native or deamidated FN for 24h. U937 cell adhesion to deamidated FN was associated with marked activation of the AP-1 signal transduction pathway but not NFκB. Each column represents the mean □ ± □ SE of three biological replicates (two technical replicates per biological replicate). Statistical differences between groups were determined by unpaired T-test.

### NGR switching to IsoDGR increases monocyte gene expression of chemotactic mediators

Since AP-1 transcription factors regulate cytokine gene expression in response to a variety of external signals,[6, 34] we next examined monocyte expression levels of pro-inflammatory mediators after 24h exposure to either native or deamidated FN. Fig. 5a and 5b show expression levels of various cytokine/chemokine genes in PMA-primed U937 and THP-1 monocytes, indicating that exposure to deamidated FN induced substantial upregulation of several chemotactic molecules including CCL2, CCL4, and IL-8, as well as the metalloprotease enzyme MMP9 and associated pro-inflammatory cytokine TNFα.

Since adhesion of monocytic cells to isoDGR-containing FN induced marked upregulation of pro-inflammatory cytokine genes, we next hypothesized that altered patterns of cytokine expression may be a key event promoting further leukocyte accumulation and establishing a ‘positive feedback loop’ that drives pathology in IHD. To test this, we established a Transwell migration assay using PMA-primed U937 cells that were seeded into the bottom chambers of culture plates that were coated with either native or deamidated FN (or uncoated control wells). After 6h incubation, unstimulated monocytes that had been pre-labelled with CFSE were seeded into the Matrigel-coated upper chambers and their migration patterns were monitored by fluorescence microscopy. As shown in Fig. 5c and 5d, U937 cells exposed to deamidated FN induced >2-fold more monocyte transmigration than did cells exposed to either native FN or uncoated control wells. These data confirmed that monocytic cell adhesion to isoDGR-containing FN triggers the release of soluble factors that promote further leukocyte recruitment to the modified matrix and likely represents a key early event in atherosclerotic plaque formation/progression.

**Fig. 5.**
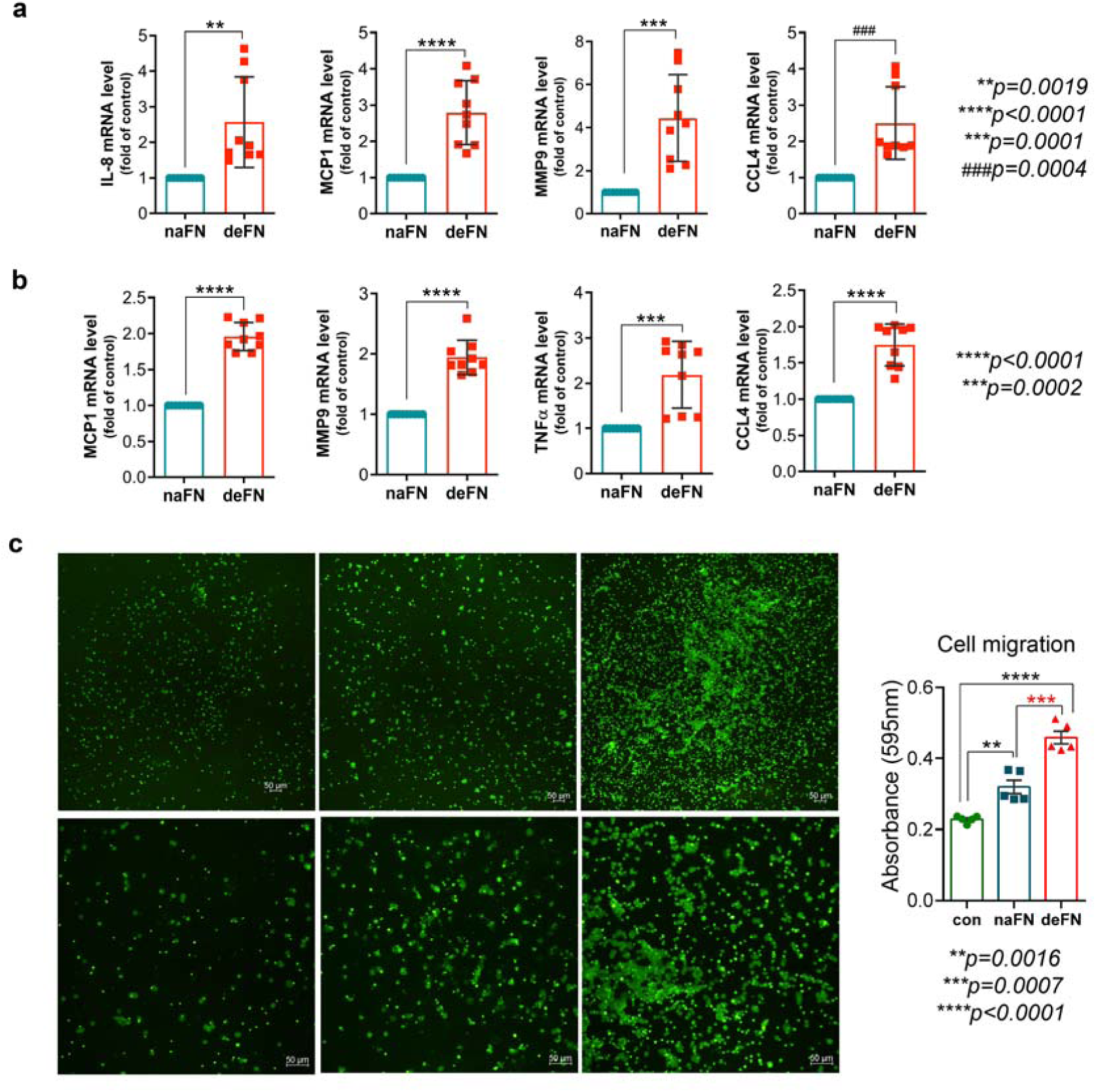
Integrin-mediated monocyte binding to FN-isoDGR promotes further leukocyte recruitment. Monocytic U937 cells (**a**) or THP-1 cells (**b**) were primed with PMA then cultured for 24h together with native or deamidated FN before mRNA expression levels of key cytokines were assessed by qRT-PCR. This analysis revealed that monocytes exposed to FN-isoDGR displayed significant upregulation of several chemotactic molecules, pro-inflammatory cytokines, and metalloproteinase enzyme MMP9. Experiments included n=3 biological replicates (3 technical replicates per biological replicate, data are shown as mean ± S.E. and were assessed by unpaired T-test). **c** In Transwell experiments, adhesion of U937 cells to deamidated FN (bottom chamber) induced greater transmigration of unstimulated monocytes (matrigel-coated top chamber) than did exposure to the native protein control (assessed by microscopic or colorimetric analysis after 24h incubation). Experiments included n=5 biological replicates. Statistical differences between groups were determined by unpaired T-test.

**Fig. 6.**
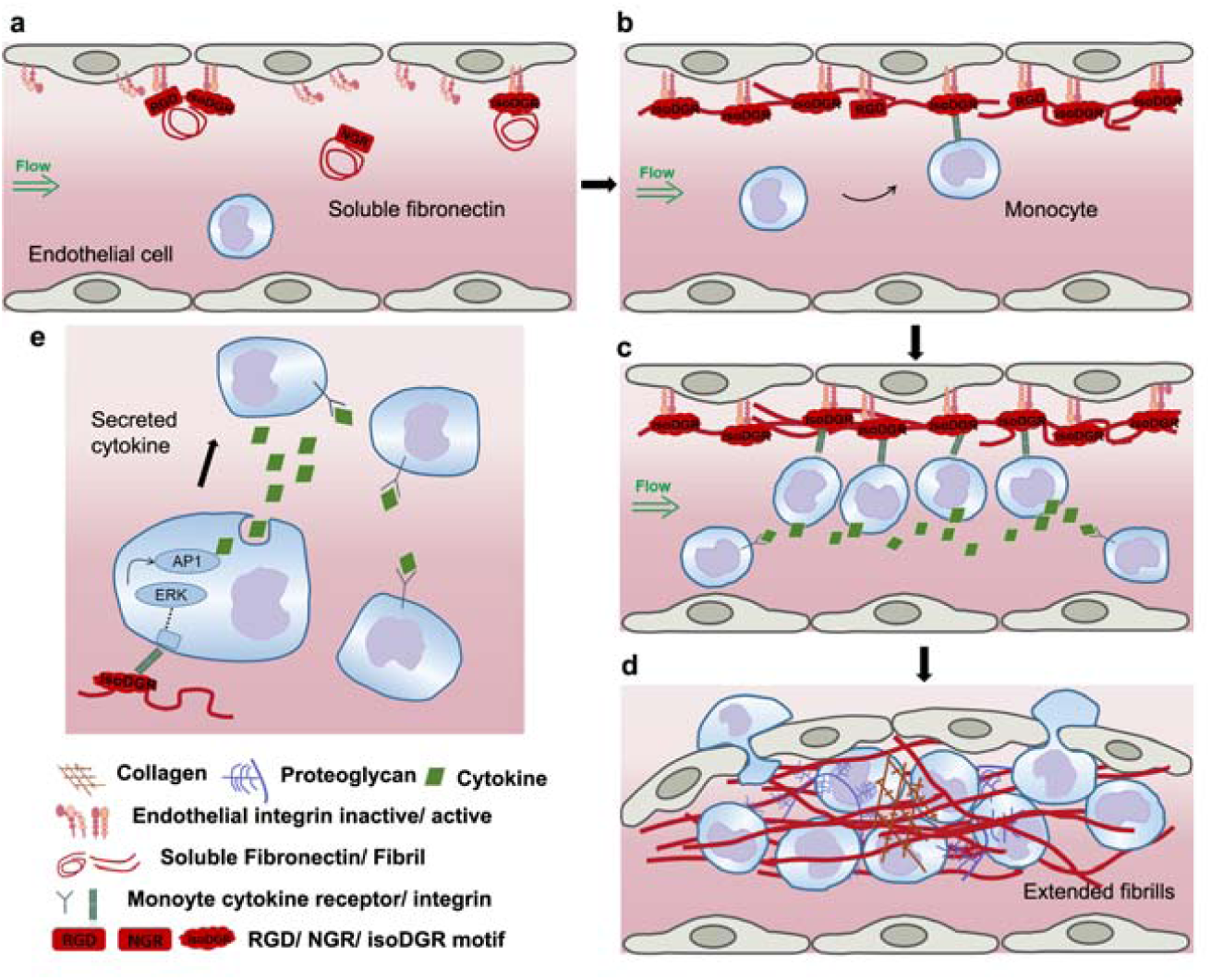
Deamidation of NGR to isoDGR in plasma fibronectin is a critical event in vascular fibrosis. **a** Accumulation of soluble plasma FN on endothelial cell surfaces is mediated by integrin-RGD or integrin-isoDGR interaction. **b** IsoDGR-containing FN accelerates protein fibrillogenesis on the surface of endothelial cells. **c** Circulating monocytes bind to FN fibril via strong interaction between monocyte integrin and isoDGR motif of FN. **d** IsoDGR-mediated integrin activation on monocytes surface promote the secretion of pro-inflammatory cytokines and further recruit the circulating leukocytes to initiate vascular fibrogenesis. **e** IsoDGR-containing FN activates the integrin:ERK:AP-1 signalling cascade to secrete the cytokines.

## Discussion

This study provides evidence that deamidation of blood plasma fibronectin (FN) creates isoDGR motifs that increase binding to β1 integrins on the surface of endothelial cells and monocytes, leading to activation of integrin ‘outside-in’ signalling and expression of cytokines that promote further leukocyte recruitment to the developing atherosclerotic matrix. These data suggest that isoDGR formation is a key early event in the pathogenesis of coronary artery disease (CAD) that can be accurately quantified using a specific monoclonal antibody-based ELISA method. It is possible therefore that similar approaches could also be used to develop routine blood tests for early detection of atherosclerosis that facilitate timely clinical intervention and reduce mortality in patients with cardiovascular disease (CVD).

CVD is a major public health issue associated with an enormous socio-economic burden as both incidence and associated mortality continue to increase around the world[35]. While many CVD-related deaths could potentially be avoided by effective heath management after early detection of key risk factors, current models of CVD prediction exhibit only limited prognostic power[1, 10, 13, 14, 22, 40]. In the current study, we therefore sought to identify critical early events in atherosclerotic plaque formation that can be detected in peripheral blood samples and could potentially be used to improve CVD risk prediction in the clinic. We have previously reported that specific matrix proteins forming carotid atherosclerotic plaques in CVD patients exhibit marked deamidation of NGR motifs[20], thus facilitating increased monocyte binding to core matrix components[16]. In the current study, we generated a novel monoclonal antibody to enable isoDGR screening of blood plasma samples, which revealed that IHD is associated with increased deamidation of fibronectin and fibrinogen proteins that are known to be critical mediators of arterial thrombosis. Together, these data led us to hypothesize that deamidated plasma proteins may regulate crosstalk between the endothelium and circulating leukocytes via effects on integrin-ligand interactions.

To address the potential role of deamidated FN in the vascular network and potential correlations with CVD pathology, we investigated the effect of isoDGR-containing FN on endothelial cell biology using a series of phenotypic and function assays *in vitro*. In these experiments, isoDGR-modified FN was consistently found to promote activation of integrin β1 in human endothelial cells due to the isoDGR motif itself serving as a potent ligand for β1 binding. Furthermore, after chemical deamidation *in vitro*, FN was capable of accumulating to high levels on endothelial cell layers where the modified protein accelerated formation of stable, insoluble fibrils under conditions of steady flow. These data strongly suggest that deamidation of NGR motifs in blood plasma FN facilitates matrix deposition on the artery intima and serves as a seed for further atherosclerotic processes. Indeed, extensive deposition of ECM is known to be a key feature of vascular fibrosis that is strongly associated with both artery stiffness and progression of atherosclerosis.[17, 26] Our observation that deamidated FN rapidly aggregates and forms extended fibrillary networks on endothelial cell layers therefore suggests that isoDGR-containing FN may contribute to enhanced vascular stiffness and reduced elasticity, likely due to the stretched/extended form of FN providing additional mechanical tension to the vessel walls[45].

Circulating monocytes can bind to ECM via integrin interactions with RGD or isoDGR motifs[12]. Since several matrix proteins are highly deamidated in atherosclerotic plaque tissues, and in particular isoDGR-containing FN readily aggregates to form fibrils, these data suggest that monocytes can readily adhere to matrix assembled on top of endothelial cells in IHD. Consistent with this concept, our data indicated that isoDGR:integrin-signalling is a specific feature of the atherosclerotic matrix that triggers differential patterns of gene expression in adherent leukocytes. Indeed, MAPK/ERK-dependent signalling and upregulation of transcription factor AP-1 were induced only in monocytic cells binding to deamidated FN, resulting in amplification of several chemokine/cytokine genes and transmigration of additional immune cells towards the assembling matrix. While AP-1 signal transduction is known to be activated by MAPK/ERK activation in response to a variety of external cues[23, 24, 34], the mechanism by which the isoDGR:integrin:ERK:AP-1 axis induces signals in CVD will require further investigation.

In summary, our data suggest that molecular switching from NGR to isoDGR motifs facilitates FN binding to integrins on the surface of endothelial cells and monocytes, leading to ‘outside-in’ signalling and induction of a range of potent pro-inflammatory mediators. Together, these events drive further leukocyte recruitment and accumulation in the arterial wall, likely aiding the progression of atherosclerosis.

## Author contribution

J.E.P designed and performed experiments, analyzed data, and wrote the manuscript; X.G. performed LC-MS/MS and ELISA analysis and wrote manuscript; K.C.K, S.S.N, S.E.L. performed qPCR and WB experiments; W.M. performed LC-MS/MS and MALDI-TOF-MS analysis; S.C.L., M.K.L., D.J.P., N.E.M., D.K., V.S, and H.H.H contributed clinical samples, reagents and revised the manuscript, S.K.S. conceived and supervised the project and revised the manuscript. All co-authors contributed to the revision of the manuscript.

## Declarations

The authors declare no conflict of interest.

