## Supplementary material for "Increased isoDGR motifs in plasma fibronectin are associated with atherosclerosis through facilitation of vascular fibrosis": Table

| | Ischemic Heart Disease (IHD) (N=25) | Control (Ctrl) (N=25) |
|---|---|---|
| Age [Mean (± SD)] | 63 (± 7.6) | 54 (± 8.7) |
| Gender [N (%)] | | |
| Male | 18(72) | 15 (60) |
| Female | 7(28) | 10(40) |
| Chinese [N (%)] | 25 (100) | 25(100) |
| Treatment [N (%)] | | |
| CABG | 25(100) | NA |
| Conservative | NA | 25(100) |
| Hypertension [N (%)] | | |
| Hypertension | 19 (76) | 5 (20) |
| No Hypertension | 6(24) | 20(80) |
| Hyperlipidemia [N (%)] | | |
| Hyperlipidemia | 25(100) | 8 (32) |
| No Hyperlipidemia | NA | 17(68) |
| Diabetes Mellitus [N (%)] | 15 (60) | 14 (70) |
| Lipid medication (statin therapy) [N (%)] | 25(100) | 10(40) |
| Smoking [N (%)] | | |
| Smoking | 9(36) | 1(4) |
| No Smoking | 14(56) | 24(96) |
| Missing information | 2(8) | NA |

N, number of individuals SD, Standard Deviation
