## Supplementary Tables for "Increased isoDGR motifs in plasma fibronectin are associated with atherosclerosis through facilitation of vascular fibrosis"

**Supplementary Table 1. Clinical characteristics of IHD patients**

| Subject No | Disease | Treatment | Gender | Ethinic | Age | EF | DM | ESRF | Smoking | Hypertension | Hyperlipidaemia | Lipid medication (statin therapy) | PVD |
|---|---|---|---|---|---|---|---|---|---|---|---|---|---|
| AE14/1471 | IHD non-MI | OPCAB | Male | Chinese | 65 | >45% | Yes | No | Yes | Yes | Yes | Yes | No |
| AE14/1475 | IHD non-MI | CABG | Male | Chinese | 68 | >45% | Yes | No | No | Yes | Yes | Yes | No |
| AE14/1486 | IHD non-MI | CABG | Male | Chinese | 70 | >45% | Yes | No | Yes | Yes | Yes | Yes | No |
| AE14/1487 | IHD non-MI | CABG | Male | Chinese | 56 | 45-35% | Yes | No | No | Yes | Yes | Yes | No |
| AE14/1488 | IHD non-MI | Redo CABG & Femoral Cannulation | Male | Chinese | 63 | >45% | No | No | | No | Yes | Yes | No |
| AE14/1505 | IHD non-MI | CABG | Male | Chinese | 53 | <35% | Yes | No | Yes | Yes | Yes | Yes | No |
| AE14/1531 | IHD non-MI | CABG | Male | Chinese | 56 | >45% | Yes | No | Yes | Yes | Yes | Yes | No |
| AE14/1534 | IHD non-MI | CABG | Male | Chinese | 71 | >45% | Yes | No | Yes | Yes | Yes | Yes | No |
| AE14/1538 | IHD non-MI | CABG | Male | Chinese | 78 | 45-35% | Yes | No | No | No | Yes | Yes | No |
| AE14/1546 | IHD non-MI | CABG | Female | Chinese | 67 | | No | No | No | Yes | Yes | Yes | No |
| AE14/1556 | IHD non-MI | CABG | Male | Chinese | 44 | 45-35% | No | No | Yes | Yes | Yes | Yes | No |
| AE14/1562 | IHD non-MI | CABG | Male | Chinese | 55 | <35% | Yes | No | No | Yes | Yes | Yes | No |
| AE14/1571 | IHD non-MI | CABG | Female | Chinese | 62 | >45% | Yes | No | No | Yes | Yes | Yes | No |
| AE14/1576 | IHD non-MI | CABG | Male | Chinese | 65 | >45% | No | No | No | Yes | Yes | Yes | No |
| AE14/1577 | IHD non-MI | CABG | Female | Chinese | 70 | >45% | Yes | No | No | Yes | Yes | Yes | No |
| AE14/1588 | IHD non-MI | CABG | Male | Chinese | 65 | >45% | No | No | Yes | Yes | Yes | Yes | No |
| AE14/1593 | IHD non-MI | CABG | Male | Chinese | 66 | >45% | Yes | No | Yes | Yes | Yes | Yes | No |
| AE14/1594 | IHD non-MI | CABG | Male | Chinese | 73 | | Yes | No | | Yes | Yes | Yes | No |
| AE14/1598 | IHD non-MI | CABG | Female | Chinese | 63 | >45% | No | No | No | Yes | Yes | Yes | No |
| AE14/1626 | IHD non-MI | Redo-sternotomy CABG | Female | Chinese | 66 | >45% | Yes | No | No | No | Yes | Yes | No |
| AE14/1638 | IHD non-MI | CABG | Male | Chinese | 65 | >45% | Yes | No | No | Yes | Yes | Yes | No |
| AE14/1647 | IHD non-MI | Minimally Invasive CABG | Male | Chinese | 55 | >45% | No | No | No | No | Yes | Yes | No |
| AE14/1681 | IHD non-MI | CABG | Female | Chinese | 67 | >45% | No | No | No | Yes | Yes | Yes | No |
| AE14/1710 | IHD non-MI | CABG | Male | Chinese | 53 | <35% | No | No | Yes | No | Yes | Yes | No |
| AE15/1827 | IHD non-MI | CABG | Female | Chinese | 60 | >45% | No | No | No | No | Yes | Yes | No |

**Supplementary Table 2. Clinical characteristics of control**

| Subject no | Group | Treatment | Gender | Race | Age | EF | DM | ESRF | Smoking | Hypertension | Hyperlipidaemia | Lipid medication (statin therapy) | PVD |
|---|---|---|---|---|---|---|---|---|---|---|---|---|---|
| SN0017 | Normal | Conservative | Male | Chinese | 31 | >45% | No | No | Yes | No | No | No | No |
| SN0040 | Normal | Conservative | Male | Chinese | 59 | >45% | No | No | No | Yes | No | No | No |
| SN0047 | Normal | Conservative | Male | Chinese | 42 | >45% | No | No | No | No | Yes | No | No |
| SN0066 | Normal | Conservative | Female | Chinese | 66 | >45% | No | No | No | No | Yes | Yes | No |
| SN0119 | Normal | Conservative | Female | Chinese | 52 | >45% | Yes | No | No | Yes | No | No | No |
| SN0122 | Normal | Conservative | Female | Chinese | 57 | >45% | No | No | No | No | Yes | Yes | No |
| SN0142 | Normal | Conservative | Male | Chinese | 48 | >45% | No | No | No | No | No | No | No |
| SN0158 | Normal | Conservative | Male | Chinese | 58 | >45% | No | No | No | No | No | No | No |
| SN0204 | Normal | Conservative | Male | Chinese | 48 | >45% | No | No | No | No | No | No | No |
| SN0211 | Normal | Conservative | Male | Chinese | 56 | >45% | No | No | No | No | Yes | Yes | No |
| SN0231 | Normal | Conservative | Male | Chinese | 47 | >45% | No | No | No | No | No | No | No |
| SN0240 | Normal | Conservative | Female | Chinese | 49 | >45% | Yes | No | No | No | No | Yes | No |
| SN0246 | Normal | Conservative | Female | Chinese | 59 | >45% | No | No | No | No | No | No | No |
| SN0271 | Normal | Conservative | Female | Chinese | 69 | >45% | No | No | No | No | Yes | Yes | No |
| SN0298 | Normal | Conservative | Male | Chinese | 47 | >45% | No | No | No | No | No | No | No |
| SN0327 | Normal | Conservative | Male | Chinese | 53 | >45% | No | No | No | Yes | No | No | No |
| SN0486 | Normal | Conservative | Male | Chinese | 64 | >45% | No | No | No | No | No | Yes | No |
| SN0518 | Normal | Conservative | Female | Chinese | 52 | >45% | No | No | No | No | Yes | Yes | No |
| SN0633 | Normal | Conservative | Male | Chinese | 65 | >45% | No | No | No | Yes | No | No | No |
| SN0665 | Normal | Conservative | Male | Chinese | 54 | >45% | No | No | No | No | No | Yes | No |
| SN0684 | Normal | Conservative | Female | Chinese | 59 | >45% | No | No | No | No | No | No | No |
| SN0788 | Normal | Conservative | Female | Chinese | 62 | >45% | No | No | No | Yes | No | Yes | No |
| SN0839 | Normal | Conservative | Male | Chinese | 46 | >45% | No | No | No | No | Yes | No | No |
| SN0846 | Normal | Conservative | Female | Chinese | 62 | >45% | No | No | No | No | Yes | Yes | No |
| SN0915 | Normal | Conservative | Male | Chinese | 61 | >45% | No | No | No | No | No | Yes | No |
